# AcrNET: Predicting Anti-CRISPR with Deep Learning

**DOI:** 10.1101/2022.04.02.486820

**Authors:** Yunxiang Li, Yumeng Wei, Sheng Xu, Qingxiong Tan, Licheng Zong, Jiuming Wang, Yixuan Wang, Jiayang Chen, Liang Hong, Yu Li

## Abstract

As an important group of proteins discovered in phages, anti-CRISPR inhibits the activity of the immune system of bacteria (*i.e*., CRISPR-Cas), showing great potential for gene editing and phage therapy. However, the prediction and discovery of anti-CRISPR are challenging for its high variability and fast evolution. Existing biological studies often depend on known CRISPR and anti-CRISPR pairs, which may not be practical considering the huge number of pairs in reality. Computational methods usually struggle with prediction performance. To tackle these issues, we propose a novel deep neural **net**work for **a**nti-**CR**ISPR analysis (**AcrNET**), which achieves impressive performance. On both the cross-fold and cross-dataset validation, our method outperforms the previous state-of-the-art methods significantly. Impressively, AcrNET improves the prediction performance by at least 15% regarding the F1 score for the cross-dataset test. Moreover, AcrNET is the first computational method to predict the detailed anti-CRISPR classes, which may help illustrate the anti-CRISPR mechanism. Taking advantage of a Transformer protein language model pre-trained on 250 million protein sequences, AcrNET overcomes the data scarcity problem. Extensive experiments and analysis suggest that Transformer model feature, evolutionary feature, and local structure feature complement each other, which indicates the critical properties of anti-CRISPR proteins. Combined with AlphaFold prediction, further motif analysis and docking experiments demonstrate that AcrNET captures the evolutionarily conserved pattern and the interaction between anti-CRISPR and the target implicitly. With the impressive prediction capability, AcrNET can serve as a valuable tool for anti-CRISPR study and new anti-CRISPR discovery, with a free webserver at https://proj.cse.cuhk.edu.hk/aihlab/AcrNET/.

## 1 INTRODUCTION

Anti-CRISPR is an important group of proteins discovered in phages for fighting against the immune system of certain bacteria. To resist the invasion of phages, bacteria have evolved different types of defense mechanisms, including the important and adaptive immune system CRISPR-Cas. Correspondingly, phages evolved inhibitor proteins Anti-CRISPRs (Acrs) to fight with the CRISPR-Cas system. Because of the strong ability to adaptively detect, destroy, and modify DNA sequences, CRISPR-Cas has become a popular geneediting tool. Since there could be a dedicated Acr available for each CRISPR-Cas system [35], performing accurate Acr predictions to find new Acrs is becoming increasingly important for many real-world applications, such as reducing off-target accidents in gene editing, measurement of gene drive, and phage therapy [26, 28, 35].

In recent years, many biological and bioinformatic approaches have been adopted to predict and discover new Acrs. Based on a plasmid-based functional assay, the activity of prophage, which integrated phage genome and bacterial genome, was observed and evaluated, leading to the discovery of the first Acr that enabled phage to replicate successfully under CRISPR-Cas attack [3]. By utilizing the BLAST search strategy on anti-CRISPR-associated (Aca) gene, which is an important characteristic of certain Acr genes, Pawluk et al. [29] developed a bioinformatic approach to find additional Acr proteins in phages and related diverse mobile genetic data in bacteria. Motivated by the idea that “self-targeting” prophage contains both a DNA target and CRISPR spacer as the CRISPR-Cas system is inactivated, Rauch et al. [32] conducted a study to search bacterial genomes for the co-existence of spacer and its target, discovering new Acrs in phages with this self-targeting phenomenon. A phage-oriented approach led to an Acr only for abolished immunity which is unrelated to previous Acrs [17]. However, these methods depend on expensive and time-consuming experimentally-generated data. Furthermore, they rely heavily on the homologs of Acrs and their functional characteristics, which is not practical for the rapid emergence of a large number of new types of proteins.

Several machine learning (ML) methods were introduced to predict Acrs and accelerate biological discoveries. An ensemble learning-based Acrs prediction tool, PaCRISPR [40], was developed to apply the support vector machine (SVM) model on evolutionary features, which were extracted from Position-Specific Scoring Matrix (PSSM), including PSSM-composition [48], DPC-PSSM [25], PSSM-AC [11], and RPSSM [8]. A new Acrs prediction model named AcRanker [12] based on XGBoost ranking was built to deal with the mixture of different sequence features, including grouped dimer, amino acid composition, and trimer frequency counts [12]. A server, AcrFinder was built to pre-screen genomic data for Acr candidates by combining three well-accepted ideas: guilt-by-association (GBA), homology search, and CRISPR-Cas self-targeting spacers [45]. A machine learning approach consisting of a random forest with 1000 trees was built to predict comprehensive Acrs, which showed strong forecasting capability for the unseen test set [13]. Also built upon the random forest, AcrDetector utilized merely six features to identify Acrs from the whole genome scale but still maintained the prediction precision [10]. As described in the AcrHub paper, it incorporates state-of-the-art predictors and three functional analysis modules, including homology network analysis, phylogenetic analysis, and similarity analysis [39]. These models are relatively flexible compared with the traditional bioinformatic approaches. However, these methods usually rely on traditional machine learning techniques, which utilize simple linear models or shallow nonlinear models, limiting their modeling capacity making it difficult to promote the development of Acr-based precise treatment. DeepAcr [38] applies deep learning models, where they only use simple protein features, lacking the ability to solve the data scarcity problem.

Considering the scarce database, Acrs’ quick evolution, and under-explored Arc features, we propose a novel deep learning approach (AcrNET) for the effective and accurate prediction and classification of Acr proteins to facilitate new Acr discovery from large-scale protein databases. The biggest challenge of developing deep learning methods to predict Acrs is the lack of data. For example, we only have 1094 non-redundant Acr sequences in our dataset, which is usually insufficient for an effective deep learning model. To deal with the data scarcity issue, we introduce a pretrained large-scale Transformer protein language model [33] to provide more informative representations. The pre-trained model extracts internal statistical properties to effectively promote the prediction performance for structure, function, and other tasks [33, 36]. This learning module explored 250 million protein sequences, significantly broadening our database and helping achieve better prediction performance. Meanwhile, this module is computationally efficient, which can even meet the time requirement used for tremendous protein sequences. Protein language models have been applied in many downstream tasks and led to improvement, including protein-protein interaction [34] and residue-residue contact prediction [31]. Furthermore, to fully utilize valuable information of protein data to promote the performance, we combine the protein language model feature with various other features, including Acr sequence information, Acr protein structure information, relative solvent accessibility, and four evolutionary features from PSSM (PSSM-composition, DPC-PSSM, PSSM-AC and RPSSM), which are proved helpful for Acrs prediction [12, 40]. We also provide more detailed and informative hierarchical classification results compared with the current classification scheme in the anti-CRISPR system, which broadly divides proteins into two categories according to whether a protein is Acr or not [21]. At the first level, similar to the current anti-CRISPR system, we predict whether a protein is an Acr. Then, at the second level, if it is an Acr protein, we further provide which class of Acrs the protein belongs to, which can bridge the large-scale protein database and Acrs to accelerate the discovery and verification of Acrs. Down to the details of prediction and classification procedure, we develop the model based on convolutional neural networks (CNNs) and deep neural networks (DNNs).

Our contributions are as follows.

- Based on the datasets from anti-CRISPRdb [9] and PaCRISPR [40], we propose a deep learning method, AcrNET, for anti-CRISPR prediction, which outperforms the previous state-of-the-art methods by at least 15% regarding the F1 score on cross-dataset validation.
- We develop the first computational method that can predict the detailed anti-CRISPR classes, which may help illustrate the anti-CRISPR mechanism.
- We are the first one in this field to resolve the data scarcity issue by transferring the knowledge from a large-scale protein language model trained on 250 million protein sequences.
- We perform extensive experiments and analysis to investigate the relation among the Transformer model feature, evolutionary feature, and local structure feature, which suggests that they complement each other, indicating the critical properties of anti-CRISPR proteins.
- Combining AcrNET with AlphaFold and motif detection methods, we propose a computational pipeline to understand the model prediction basis better and validate our prediction computationally.

## 2 METHOD

### Overview of AcrNET

AcrNET contains three basic parts, as in Fig. 1. First, a large collection of protein sequences are compiled into the pipeline to extract protein features, including secondary structure, relative solvent accessibility, four evolutionary features, and Transformer features. Furthermore, by introducing the Transformer learning module, we tackle the scarcity issue of Acrs database with a large-scale pre-training database. Second, the learned features are injected into AcrNET, which jointly models these features using two deep learning modules, CNN and DNN, in an end-to-end trainable deep architecture. To this end, the model provides the confidence score of the likelihood that the protein is an Acr protein. Finally, based on the predicted likelihood, we can perform some downstream tasks. For example, positive Acr candidates are further inputted into AcrNET to perform the second-level prediction, predicting which sub-category the Acr belongs to. This sub-category prediction helps narrow down the candidate list, assisting biologists in carrying out biological experiments. We also conduct motif analysis to illustrate the implicit features learned by AcrNET. Additionally, we show the interaction details between the Acr protein and the target by predicting structure with AlphaFold and conducting protein-protein docking.

**Figure 1:**
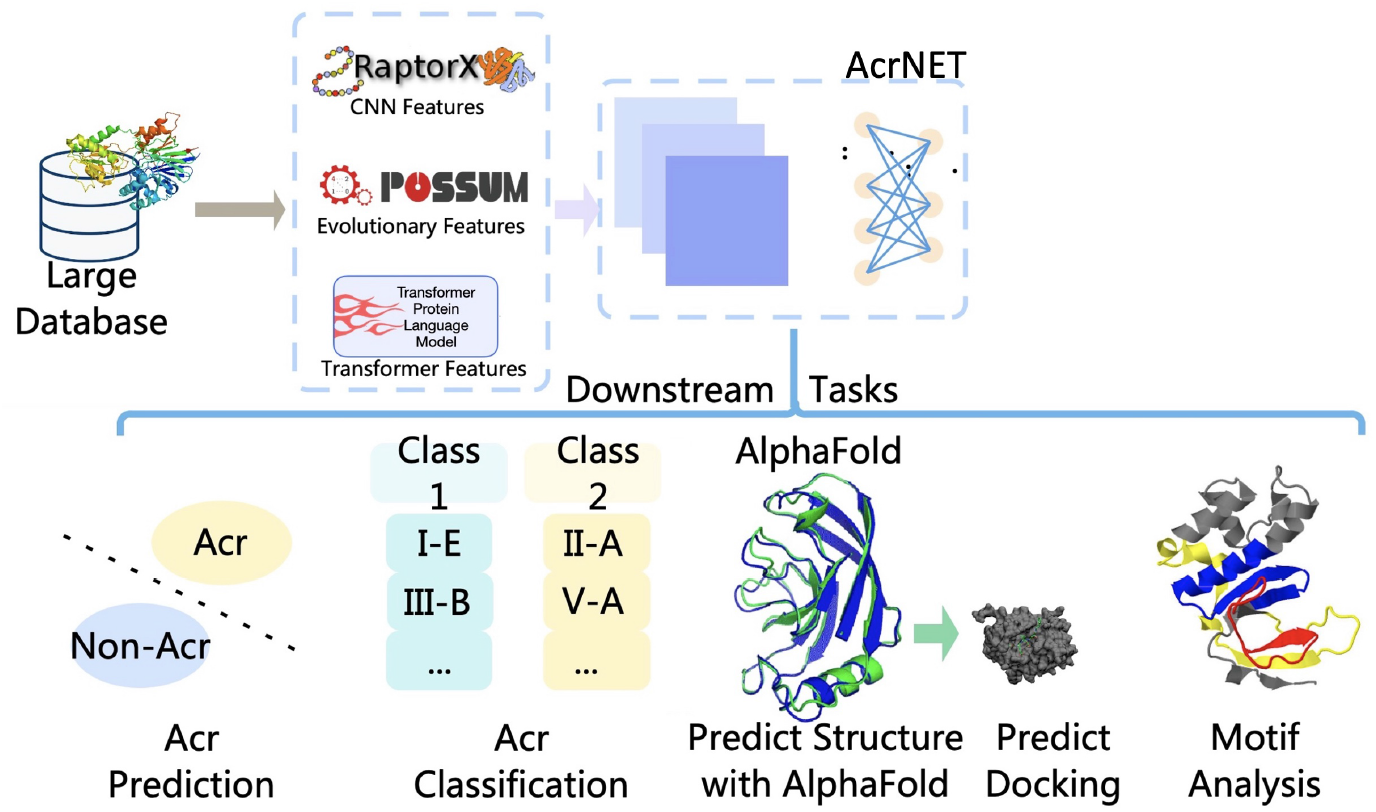
The AcrNET pipeline. The process contains three steps. Firstly, we compile the large protein sequence database to obtain secondary structure, relative solvent accessibility, evolutionary features, and Transformer features. Secondly, based on the features, the AcrNET model predicts the Acr proteins. Finally, with the predicted likelihood of the protein being an Acr protein, we can perform multiple downstream tasks, including predicting classes of Acrs, doing biological validation experiments, predicting the Acr structure with AlphaFold, investigating the interaction between Acr and the target with protein-protein docking, and motif analysis.

### Extracting evolutionary features and structure features for comprehensive protein representation

Protein features necessary for Acr prediction are not well-defined. Thus, we focused on evolutionary and sequence features that have proven useful in earlier studies [39, 40]. We also incorporated the secondary structure, which is applied in protein function prediction [24, 49] and known to be related to protein-protein interaction [18]. First, we use one-hot encoding to encode the protein sequence, which contains original amino acid information, reflecting vital sequential patterns of Acr proteins. Second, we extract PSSM features to introduce evolutionary information of proteins. The PSSM score is an *L* × 20 matrix summarizing the homologs similar to a given protein sequence in the specified database. Each score in the matrix reflects the conservation of the corresponding amino acid at this position, where large scores imply high conservation. From the PSSM matrix, we calculate four evolutionary features, PSSM-composition, DPC-PSSM, PSSM-AC, and RPSSM, which are fixed-size matrices. They deal with PSSM varying lengths, encode relation between two elements, and local sequence order effect [40]. Third, we extract secondary structure information with traditional three classes [27] and eight classes extended by Kabsch and Sander [19], and convert them with one-hot encoding. We also consider solvent accessibility which presents the local exposure of a protein. We label them with three states and apply one-hot encoding. The last type of feature that we consider is Transformer feature, which helps consider the biological structure and function of protein data. During the process of learning such features, we can simultaneously handle the scarcity issue of Acrs database. We discuss it as a separate section in the following paragraph.

### Introducing Transformer learning to tackle data scarcity

The proposed deep learning model has a large number of trainable parameters, which enables the model to have a very strong modeling capacity. However, it may suffer from the overfitting problem for data scarcity issue, especially for Acr prediction, where we only have around 1000 sequences. Thus we introduce the Transformer learning algorithm to learn more informative representations [33], significantly broadening the training database. Unsupervised method has been introduced with the explosive growth of protein sequence data, which can capture statistical regularities [14]. This module learns the sequential information of the protein data by predicting the contents of the masked parts, therefore guiding it to learn useful structure information from the sequential data to provide effective representations. Specifically, the objective function is designed as follows to minimize the masked language modeling (MLM) loss:

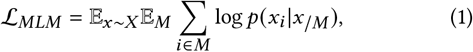

where *x* represents a protein sequence; *M* is the set of mask indices; *x_i_* denotes a protein sequence with mask token at index *i*. Furthermore, because when we train the language model, we do not need the human-labeled data, we can train the model with as many protein sequences as possible. We adopt a protein language model trained over 250 million protein sequences from UniProt. The authors of ESM-1b explored datasets with up to 250 million sequences of the UniParc database [2], which has 86 billion amino acids. The dataset they used owns comparable size to large text datasets that are being used to train high-capacity neural network architectures on natural [7, 30]. They trained ESM-1b on the high-diversity sparse dataset (UR50/S) uses the UniRef50 representative sequence. With such a huge amount of unsupervised training data, the model can implicitly learn the data distribution, evolutionary information, and protein sequential structure, leading to an effective protein representation. In addition, compared with the time and memory-consuming PSI-BLAST, POSSUM [41] and RaptorX [20], the introduced ESM-1b module [4, 46] is able to generate Transformer features much more quickly and more simply.

### Input data

We collect the anti-CRISPRs samples and non-anti-CRISPRs samples from Anti-CRISPRdb [9] and PaCRISPR [40], respectively. We firstly describe the process of collecting non-redundant positive samples from Anti-CRISPRdb [9]. More than 3000 experimentally characterized Acrs are initially collected [9] and CD-HIT is then applied to these samples to remove the redundant sequences by using a 95% identity threshold. After this selection process, we obtain 1094 non-redundant protein sequences of the Acrs. Next, we introduce the procedures of constructing the negative training dataset and the cross-dataset test dataset from PaCRISPR [40]. The non-anti-CRISPRs samples in this dataset [40] are all from Acr-containing phages or bacterial mobile genetic elements. The sequence sizes of these samples range from 50 to 350, with the inter-similarity of less than 40%. By using the criteria for selecting the negative sample proteins as introduced in Wang et al. [40], we obtain 902 negative samples from the negative training dataset and 260 negative samples from the negative testing dataset for cross-dataset test. The 1094 non-redundant positive samples from Anti-CRISPRdb, the 902 negative samples from the negative training dataset, and 260 negative samples from the cross-dataset dataset of Wang et al. [40], are combined as the whole experiments dataset, where the sequence similarity between positive samples and negative samples is also less than 40%. Since the sequence similarity between the training samples and the test samples is less than 40%, this cross-dataset test can effectively reflect the generalization ability of the proposed model. After data collection, we appled POSSUM, RaptorX and ESM-1b to obtain corresponding features as we mentioned in **Extracting evolutionary features and structure features for comprehensive protein representation.** session.

To collect the dataset for the task of the Acr type prediction task, we utilize the non-redundant samples in Anti-CRISPRdb, which contains 12 types of Acrs. The names of these 12 types of Acrs and the corresponding number of collections for each kind are in Table 1. In our design, we re-group these 12 types into 5 types. Specifically, the samples in the first four types, namely II-A, I-F, I-D, and II-C, are maintained, while the rest 8 types with very few samples are grouped into the ‘fifth type’. Therefore, as described in the Results section, the Acrs classification problem is a 5-class classification task.

**Table 1:** Statistical summary of the Acr classes in the non-redundant Anti-CRISPRdb dataset. We keep the largest four classes and group the rest eight smaller classes as class five.

| Acr type | II-A | I-F | I-D | II-C | I-E | V-A | I-C | VI-A | VI-B | III-1 | III-B | I-B |
| --- | --- | --- | --- | --- | --- | --- | --- | --- | --- | --- | --- | --- |
| Collection | 828 | 134 | 46 | 30 | 19 | 15 | 8 | 7 | 7 | 1 | 1 | 1 |

### Evaluation test setup

To evaluate the performance of our deep learning model, we adopt 5-fold cross-validation tests on the dataset, consisting of 1094 positive samples and 1162 negative samples, as described in the previous section. Furthermore, another cross-dataset test is conducted to evaluate the generalization ability of the proposed model for the Acr prediction. In the cross-dataset test, the Acrs of types I-F, II-C, and I-D are selected as the positive test samples, and the rest Acrs are used as the positive training samples for separation 1. I-F, I-E, V-A, I-C, VI-A, VI-B, III-I, III-B, and I-B are selected as the positive test samples for separation 2, and I-D, II-C, I-E, V-A, I-C, VI-A, VI-B, III-I, III-B, and I-B are selected as the positive test samples for separation 3. Since the sequence similarity between the training samples and the test samples is less than 40%, this cross-dataset test can effectively reflect the generalization ability of the proposed model. For negative samples, we use the separation method provided by PaCRISPR to separate samples, which means 902 negative samples from the negative training dataset of PaCRISPR are used as negative training samples, while 260 negative samples from the cross-dataset dataset of PaCRISPR are used as negative test samples. The specific arrangements are illustrated in Table 2.

**Table 2:** Statistics summary of the dataset used in our experiments. We combine Anti-CRISPRdb dataset and PaCRISPR dataset, and remove the sequence redundancy.

|  | Cross-dataset training | Cross-dataset testing |
| --- | --- | --- |
| Positive (separation 1) | 884 | 210 (From type I-F, II-C, and I-D in anti-CRISPRdb) |
| Positive (separation 2) | 904 | 190 (From type I-F, I-E, V-A, I-C, VI-A, VI-B, III-I, III-B, and I-B in anti-CRISPRdb) |
| Positive (separation 3) | 962 | 132 (From type I-D, II-C, I-E, V-A, I-C, VI-A, VI-B, III-I, III-B, and I-B in anti-CRISPRdb) |
| Negative | 902 | 260 |

### Architecture for complex feature learning and Acr prediction and classification

Since original information from proteins is complex, we leverage CNNs and DNNs to learn useful features, designed in the end-to-end trainable way as illustrated in Fig. 2. The one-hot matrices of sequence, structure feature, and solvent accessibility are concatenated together as the inputs of the CNN module. The convolutional layers learn deep features by extracting important local information from the one-hot matrices. After this, the max-pooling layer is attached along the sequential direction to calculate the largest value in each feature map. The features learned by the CNN module are further combined with the evolutionary features and Transformer features from ESM-1b, which are then jointly modeled by the DNN module. The Acr prediction is a binary classification task, which makes an estimation about whether a certain protein sequence is an Acr. Therefore, the final outputs are produced by fully connected layers with a two-unit output.

**Figure 2:**
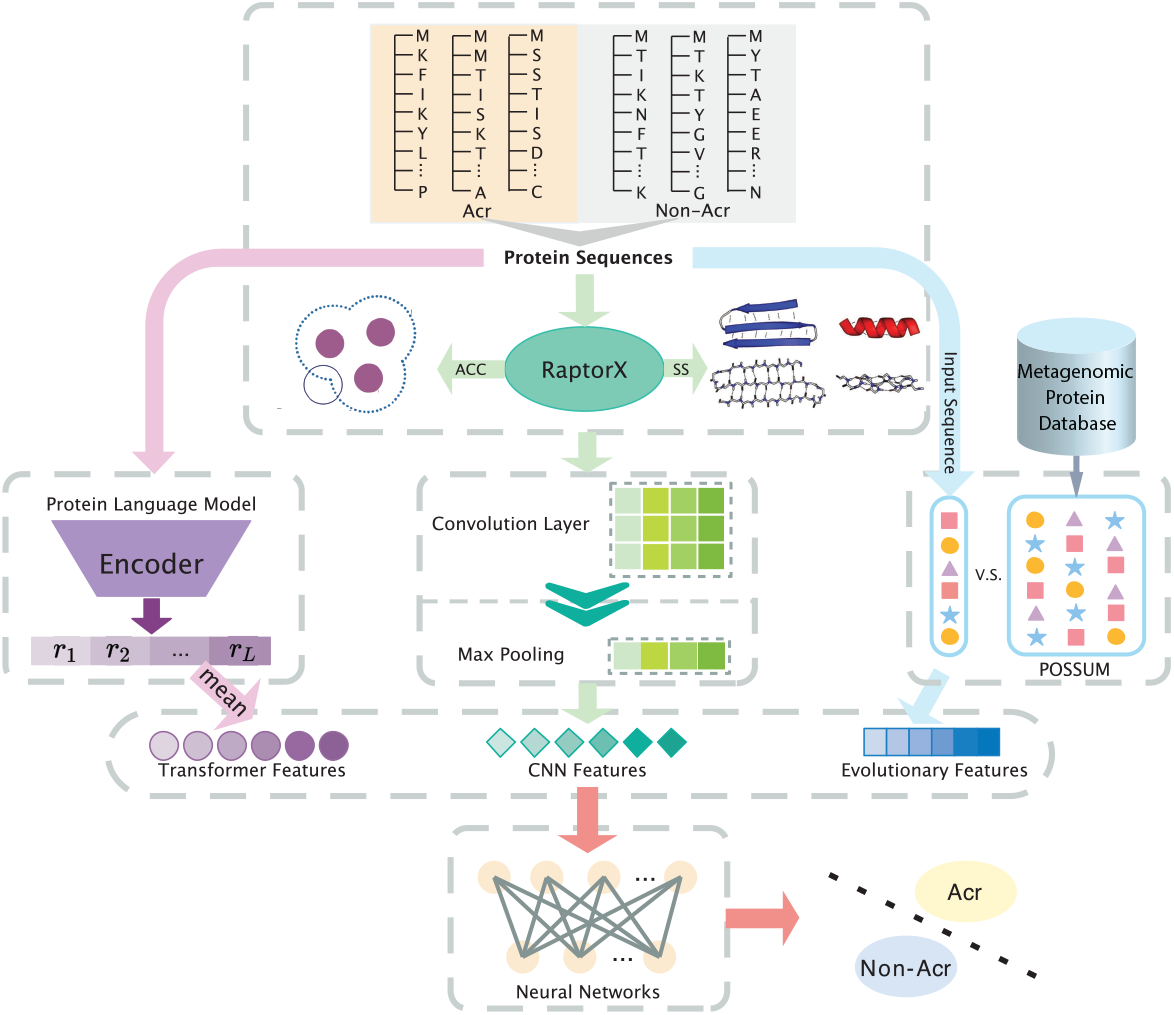
The architecture of the proposed AcrNET model. We first obtain the relative solvent accessibility, secondary structure, evolutionary features, and Transformer features with RaptorX, POSSUM, and a pretrained protein language model. Then, we concatenate the features and input them to CNNs and DNNs to obtain the predicted likelihood of the protein being an Acr protein.

Considering biologists may also be interested in Acr types, if a protein is predicted to be an Acr, we will further estimate which kind of Acr this protein belongs to and provide more detailed information for the downstream biological experimental verification. Due to the limited number of Acr samples, we separate this task from Acr prediction to avoid the imbalanced data issue and defined it as the Acr classification task, predicting the specific category of the Acr. This classification task utilizes a similar structure to the Acr prediction task with slight modification, namely changing the dimension of the final output from two to five classes. In addition, we remove the non-anti-CRISPRs samples from the whole samples and only analyze samples predicted to be Acr proteins. We use cross-entropy as the loss function and Adam as the optimizer to train the model. Moreover, to increase the flexibility of the model for real-world applications, we make the model accept the input with variant features, which allows users to select the input features according to their needs.

### Anti-CRISPR prediction performance

In this section, we discuss the performance of our model on predicting whether a certain protein sequence is an Acr or not. To sufficiently compare the prediction performance of our proposed method and existing models, we perform five-fold cross-validation test and an additional crossdataset test to evaluate their generalization ability. To exclude the influence of random seeds, we randomly generate seeds ten times for the initialization of model parameters and the split of samples of training and test groups. The final result of each model is their average of ten times prediction results. Five evaluation metrics, namely accuracy (ACC), precision, recall, F1-score, and Matthews correlation coefficient (MCC), are utilized to evaluate the prediction results. We also report TP (true positive), FN (false negative), FP (false positive), TP (true positive) and specificity for ambiguous performance for further analysis. Since other methods require additional information other than the sequence alone, such as gene location on chromosome [10], we compare AcrNET with three recently proposed methods with the same dataset we collected, namely PaCRISPR [40], AcRanker [12], and DeepAcr [38]. All of them have shown strong Acr predicting capacities and efficiency.

The results of five-fold cross-validation test and the cross-dataset test on different separations are demonstrated in Table 3, Table 4, Table 5 and Table 6 respectively, which clearly indicates that our proposed method significantly outperforms PaCRISPR, AcRanker and DeepAcr. Some important discoveries can be observed from these results. First, AcrNET substantially outperforms other methods in both tests, especially in the cross-dataset test. With respect to TP and TN, AcrNET with only Transformer features and the complete AcrNET are better on both TP and TN. The *p*-values for AcrNET performing better than PaCRISPR, AcRanker and DeepAcr are all < 0.0001. In the five-fold cross-validation test, the F1 score is promoted by around 10%. The ROC curves also suggest that AcrNET is more robust than the other methods (Fig. 3(C)). Significantly, AcrNET achieves at least 15% improvement regarding F1 score in the cross-dataset test, where training and testing data have different distributions. The results demonstrate that AcrNET is more suitable for the real Acr prediction task. We assume that AcrNET method can automatically learn more useful knowledge from the protein data, including the important structure information, which enables AcrNET to outperform other models. Furthermore, the Transformer model that we introduce to our model contains common properties of protein sequences from an extensive database, which effectively improve the generalization ability for unseen sequences.

**Figure 3:**
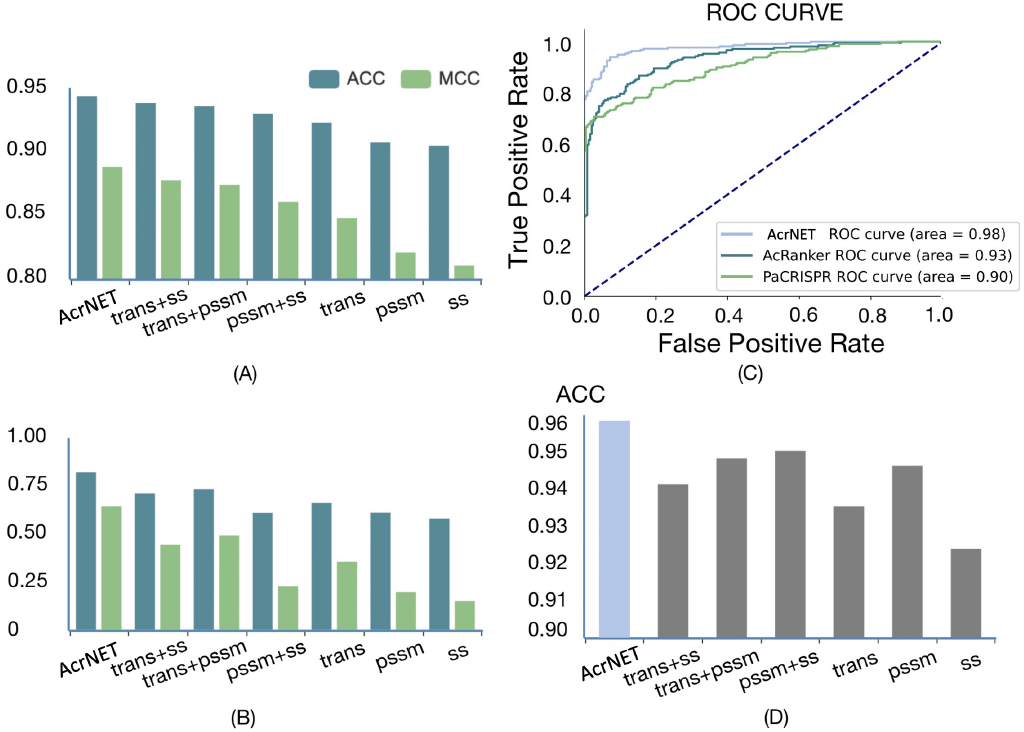
Feature influence on the prediction results. “ss” represents sequence encoding, 3-class secondary structure, 8-class secondary structure, and solvent accessibility; “trans” indicates Transformer features; “pssm” denotes 4 evolutionary features extracted from PSSM. Results in the figure are averaged over 10 different random seeds in our experiments. (A) Contribution evaluation of different features on five-fold cross-validation test. (B) Contribution evaluation of different features on cross-dataset test. (C) Performance comparison on Acr prediction regarding ROC curves of AcrNET, AcRanker and PaCRISPR. (D) Contribution evaluation of different features for the detailed classification task.

**Table 3:** Five-fold cross-validation test results of anti-CRISPRs prediction. AcrNET achieves the best results, outperforming the other methods significantly. AcrNET model with only Transformer features also has relatively good performance, better than PaCRISPR, AcRanker and DeepAcr. Results in this table are averaged over 10 different random seeds in our experiments, and variances are smaller than 0.001.

| Metrics | Accuracy | Precision | Recall | F1 score | MCC |
| --- | --- | --- | --- | --- | --- |
| AcRanker | 0.8752 | 0.8591 | 0.8953 | 0.8666 | 0.7509 |
| PaCRISPR | 0.8087 | 0.7344 | **0.9738** | 0.7588 | 0.6552 |
| DeepAcr | 0.7269 | 0.9447 | 0.6518 | 0.7710 | 0.5093 |
| AcrNET (Transformer only) | 0.9326 | 0.9351 | 0.9248 | 0.9299 | 0.8652 |
| **AcrNET** | **0.9442** | **0.9471** | 0.9409 | **0.9418** | **0.8883** |

**Table 4:** Cross-dataset test results of anti-CRISPRs prediction with separation 1. AcrNET accomplishes great improvement compared with the other state-of-the-art computational methods, especially on precision, F1 score, and MCC. The Transformer features contribute greatly to the AcrNET generalization capability, but the other features are also very helpful. Results in this table are averaged over 10 different random seeds in our experiments, and variances are smaller than 0.001.

| Metrics | TN | FN | FP | TP | Specificity | Accuracy | Precision | Recall | F1 score | MCC |
| --- | --- | --- | --- | --- | --- | --- | --- | --- | --- | --- |
| AcRanker | 244 | 157 | 16 | 53 | 0.9385 | 0.7681 | 0.2524 | 0.6319 | 0.3799 | 0.2681 |
| PaCRISPR | 217 | 67 | 43 | 143 | 0.8346 | 0.7660 | 0.7688 | **0.6810** | 0.7222 | 0.5242 |
| DeepAcr | 174 | 73 | 86 | 137 | 0.6692 | 0.6617 | 0.6143 | 0.6524 | 0.6328 | 0.3202 |
| AcrNET (Transformer only) | 255 | 110 | 5 | 100 | **0.9808** | **0.9524** | 0.4762 | 0.7553 | 0.6349 | 0.5454 |
| **AcrNET** | 232 | 57 | 28 | 143 | 0.8923 | 0.7979 | **0.8363** | **0.6810** | **0.7505** | **0.5924** |

**Table 5:**
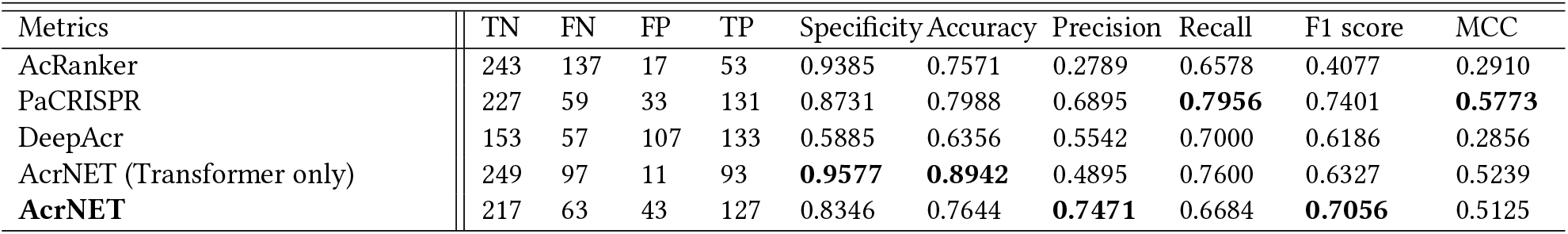
Cross-dataset test results of anti-CRISPRs prediction with separation 2.

**Table 6:** Cross-dataset test results of anti-CRISPRs prediction with separation 3.

| Metrics | TN | FN | FP | TP | Specificity | Accuracy | Precision | Recall | F1 score | MCC |
| --- | --- | --- | --- | --- | --- | --- | --- | --- | --- | --- |
| AcRanker | 238 | 109 | 22 | 23 | 0.9154 | 0.5111 | 0.1742 | 0.6658 | 0.2599 | 0.1329 |
| PaCRISPR | 224 | 32 | 36 | 100 | 0.8615 | 0.7353 | **0.7576** | 0.8265 | **0.7463** | **0.6147** |
| DeepAcr | 166 | 68 | 94 | 64 | 0.6385 | 0.5867 | 0.4051 | 0.4848 | 0.4414 | 0.1188 |
| AcrNET (Transformer only) | 248 | 91 | 12 | 41 | **0.9538** | **0.7736** | 0.3106 | 0.7372 | 0.4432 | 0.3655 |
| **AcrNET** | 162 | 8 | 98 | 124 | 0.6231 | 0.7296 | 0.5586 | **0.9394** | 0.7006 | 0.5364 |

Cross-dataset test evaluates the model’s generalization capacity when dealing with testing data with low sequence similarity from training data. From the results of the cross-dataset test, we can observe that both AcRanker and DeepAcr achieve small Precision and F1-score scores, which may be because they tend to predict new sequences as positive and cause wrong predictions. However, our model can effectively avoid the wrong predictions because the rich knowledge about the extensive database contained in the Transformer learner can help AcrNET make better predictions based on global considerations.

We also compare AcrNET with Gussow et al. [13]. The dataset used for the evaluation was collected from the study of Gussow et al. [13]. There are 3654 Acrs and 4500 non-Acrs in the dataset, identified by unique identifiers. To generate the sequences of these proteins, we searched the NCBI protein database with these identifiers using Entrez (https://www.ncbi.nlm.nih.gov/sites/batchentrez). We found that some identifiers are defined manually and cannot be found by the program. Finally, we obtained 488 positive sequences and sampled 488 balanced negative sequences. We compared the AcRanker, PaCRISPR, DeepAcr, Gussow et al. [13] and AcrNET with the same dataset. The results are listed in Table 7. We can see AcrNET outperforms all the other methods across different metrics.

**Table 7:** Five-fold cross-validation test results of Gussow et al. [13] and AcrNET. We collected training and testing data from Gussow et al. [13], used identifiers to fetch sequence data. We finally obtained 488 positive sequences and sampled balanced 488 negative sequences. We compare the previous works and AcrNET with the dataset from Gussow et al. [13].

| Metrics | Accuracy | Precision | Recall | F1 score | MCC |
| --- | --- | --- | --- | --- | --- |
| AcRanker | 0.9286 | 0.9238 | 0.9324 | 0.9276 | 0.8572 |
| PaCRISPR | 0.9367 | 0.9847 | 0.8920 | 0.9356 | 0.8779 |
| DeepAcr | 0.6796 | **0.9867** | 0.6122 | 0.7553 | 0.4521 |
| Gussow et al. [13] | 0.8493 | 0.8913 | 0.8186 | 0.8531 | 0.6992 |
| **AcrNET** | **0.9543** | 0.9762 | **0.9360** | **0.9553** | **0.9101** |

### Prediction performance on datasets with different similarity

95% similarity threshold of training dataset might be too high in practical experiments. We picked samples with 40% and 70% similarity from the positive dataset, which left 238 positive samples with the 40% threshold and 657 samples with the 70% threshold. The similarity among the negative samples was already below 40%. In the five-fold test, we randomly picked 20% samples for testing. And in cross-dataset test, the testing data from the original dataset remained the testing data, which contained 85 samples in the 40% threshold and 160 samples in the 70% threshold in separation 1, 88 samples in the 40% threshold and 156 samples in 70% threshold in separation 2, and 69 samples in 40% threshold and 108 samples in the 70% threshold in separation 3. We randomly sampled the same size of the positive dataset from the negative dataset for training and testing in 40% and 70% threshold experiments correspondingly. With the same dataset, we compared the performance between AcRanker, PaCRISPR, DeepAcr and AcrNET. The results are listed in Table 8 to Table 15. As we can see from the tables, AcrNET outperforms AcRanker, PaCRISPR and DeepAcr, especially in cross-data experiments. Analyzing the TP and TN, we can conclude that AcrNET is relatively table on TP and outperforms on TN. The performance is consistent with 95% similarity dataset experiments. We can see AcRanker performs similarly well on TN and TP, while the model and features are not powerful enough to capture more characteristics of Ars. PaCRISPR is relatively robust on TP and TN, whereas not as satisfying as ActNET, since AcrNET considers more features than PaCRISPR. And DeepAcr can be further improved on TN, we reckon that the features that DeepAcr includes may vary on non-anti-CRISPR proteins, which makes the model confused to extract common features.

**Table 8:**
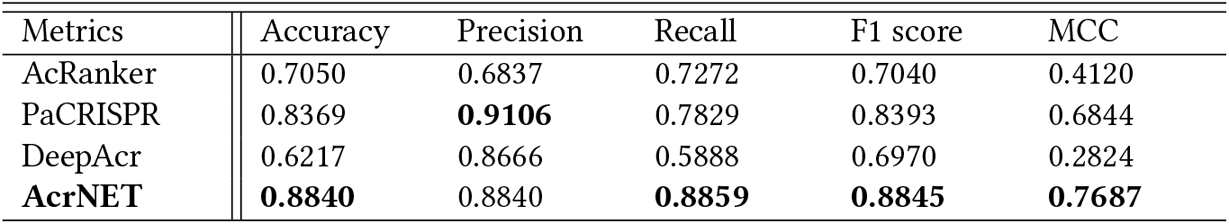
Five-fold cross-validation test results with 40% similarity dataset. We used dataset with 40% and 70% similarity and compared the performance. We used the same training and testing dataset split as previous experiments. AcrNET also outperforms other methods, the results are consistent with the previous performance.

| Metrics | Accuracy | Precision | Recall | F1 score | MCC |
| --- | --- | --- | --- | --- | --- |
| AcRanker | 0.7050 | 0.6837 | 0.7272 | 0.7040 | 0.4120 |
| PaCRISPR | 0.8369 | **0.9106** | 0.7829 | 0.8393 | 0.6844 |
| DeepAcr | 0.6217 | 0.8666 | 0.5888 | 0.6970 | 0.2824 |
| **AcrNET** | **0.8840** | 0.8840 | **0.8859** | **0.8845** | **0.7687** |

**Table 9:** Cross-dataset test results with 40% similarity dataset and separation 1.

| Metrics | TN | FN | FP | TP | Specificity | Accuracy | Precision | Recall | F1 score | MCC |
| --- | --- | --- | --- | --- | --- | --- | --- | --- | --- | --- |
| AcRanker | 58 | 21 | 27 | 64 | 0.6824 | 0.7033 | 0.7529 | 0.7176 | 0.7273 | 0.4364 |
| PaCRISPR | 66 | 15 | 19 | 70 | 0.7765 | **0.7865** | 0.8235 | 0.8000 | **0.8046** | **0.6007** |
| DeepAcr | 24 | 11 | 61 | 74 | 0.2824 | 0.5765 | 0.5481 | **0.8706** | 0.6727 | 0.1891 |
| **AcrNET** | 77 | 30 | 8 | 55 | **0.9059** | 0.7765 | **0.8730** | 0.6471 | 0.7432 | 0.5724 |

**Table 10:** Cross-dataset test results with 40% similarity dataset and separation 2.

| Metrics | TN | FN | FP | TP | Specificity | Accuracy | Precision | Recall | F1 score | MCC |
| --- | --- | --- | --- | --- | --- | --- | --- | --- | --- | --- |
| AcRanker | 69 | 35 | 19 | 53 | 0.7841 | 0.7361 | 0.6023 | 0.6932 | 0.6625 | 0.3929 |
| PaCRISPR | 69 | 33 | 19 | 55 | 0.7841 | 0.7432 | 0.6250 | 0.7045 | 0.6790 | 0.4144 |
| DeepAcr | 42 | 27 | 46 | 61 | 0.4772 | 0.5852 | 0.5701 | 0.6932 | 0.6256 | 0.1746 |
| **AcrNET** | 76 | 17 | 12 | 71 | **0.8636** | **0.8352** | **0.8554** | **0.8068** | **0.8304** | **0.6715** |

**Table 11:**
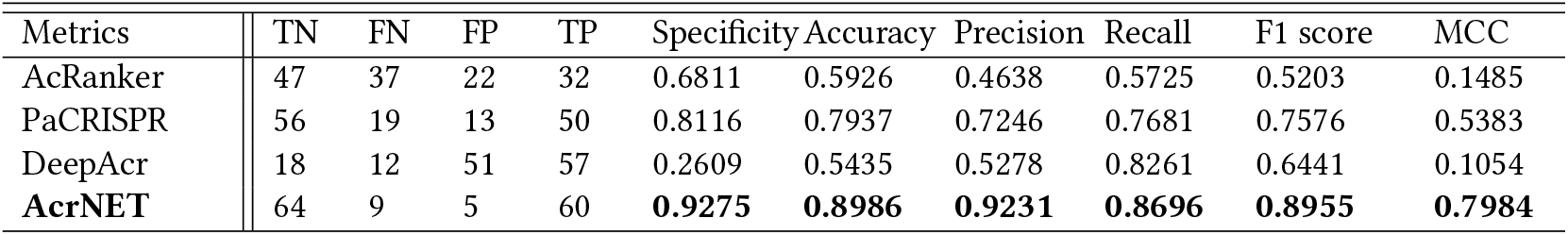
Cross-dataset test results with 40% similarity dataset and separation 3.

**Table 12:** Five-fold cross-validation test results with 70% similarity dataset.

| Metrics | Accuracy | Precision | Recall | F1 score | MCC |
| --- | --- | --- | --- | --- | --- |
| AcRanker | 0.8560 | 0.8228 | 0.8730 | 0.8466 | 0.7126 |
| PaCRISPR | 0.8799 | **0.9788** | 0.8703 | 0.9212 | 0.6933 |
| DeepAcr | 0.6211 | 0.9171 | 0.5798 | 0.7090 | 0.2965 |
| **AcrNET** | **0.9280** | 0.9480 | **0.9117** | **0.9292** | **0.8572** |

**Table 13:** Cross-dataset test results with 70% similarity dataset and separation 1.

| Metrics | TN | FN | FP | TP | Specificity | Accuracy | Precision | Recall | F1 score | MCC |
| --- | --- | --- | --- | --- | --- | --- | --- | --- | --- | --- |
| AcRanker | 135 | 88 | 25 | 72 | 0.8438 | **0.7423** | 0.4500 | 0.6469 | 0.5603 | 0.3196 |
| PaCRISPR | 128 | 105 | 32 | 55 | 0.8000 | 0.6322 | 0.3438 | 0.5719 | 0.4453 | 0.1615 |
| DeepAcr | 53 | 43 | 107 | 117 | 0.3313 | 0.5312 | 0.5223 | **0.7312** | 0.6094 | 0.0682 |
| **AcrNET** | 155 | 84 | 5 | 76 | **0.9688** | 0.7219 | **0.9383** | 0.4750 | **0.6307** | **0.5103** |

**Table 14:**
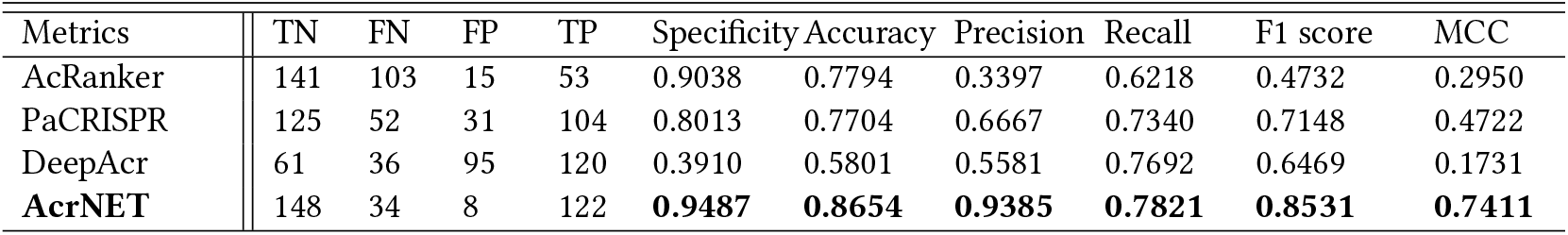
Cross-dataset test results with 70% similarity dataset and separation 2.

| Metrics | TN | FN | FP | TP | Specificity | Accuracy | Precision | Recall | F1 score | MCC |
| --- | --- | --- | --- | --- | --- | --- | --- | --- | --- | --- |
| AcRanker | 141 | 103 | 15 | 53 | 0.9038 | 0.7794 | 0.3397 | 0.6218 | 0.4732 | 0.2950 |
| PaCRISPR | 125 | 52 | 31 | 104 | 0.8013 | 0.7704 | 0.6667 | 0.7340 | 0.7148 | 0.4722 |
| DeepAcr | 61 | 36 | 95 | 120 | 0.3910 | 0.5801 | 0.5581 | 0.7692 | 0.6469 | 0.1731 |
| **AcrNET** | 148 | 34 | 8 | 122 | **0.9487** | **0.8654** | **0.9385** | **0.7821** | **0.8531** | **0.7411** |

**Table 15:** Cross-dataset test results with 70% similarity dataset and separation 3.

| Metrics | TN | FN | FP | TP | Specificity | Accuracy | Precision | Recall | F1 score | MCC |
| --- | --- | --- | --- | --- | --- | --- | --- | --- | --- | --- |
| AcRanker | 96 | 79 | 12 | 29 | 0.8889 | 0.7073 | 0.2685 | 0.5787 | 0.3893 | 0.2007 |
| PaCRISPR | 84 | 31 | 24 | 77 | 0.7778 | 0.7624 | 0.7130 | **0.7454** | 0.7368 | 0.4918 |
| DeepAcr | 53 | 35 | 55 | 73 | 0.4907 | 0.5833 | 0.5703 | 0.6759 | 0.6186 | 0.1696 |
| **AcrNET** | 102 | 36 | 6 | 72 | **0.9444** | **0.8056** | **0.9231** | 0.6667 | **0.7742** | **0.6361** |

### Evaluation of feature influence on prediction results

We perform ablation studies on different features with the five-fold cross-validation and cross-dataset tests with the full dataset to evaluate their influence on prediction results, as shown in Fig. 3 (A) and (B). We can draw some conclusions from this study. Firstly, the performances of the combination of features are always better than the performances of features alone, which means the three kinds of features complement each other. They all reflect the different aspects of the Acr properties. Secondly, Transformer features play critical roles in both five-fold cross-validation and cross-dataset tests. Especially, Transformer features always lead to the best results among all the features and meanwhile can improve the results more significantly than other features. This indicates that the Transformer learning module can effectively learn valuable knowledge from the large database to promote the Acr prediction. Thirdly, due to the task’s difficulty, the structure information prediction would not be 100% accurate, which may influence the final prediction performance and thus should be taken into consideration when using it for prediction. Relatively, evolutionary features and Transformer features are more reliable, which can always promote the downstream tasks, such as Acr prediction. Lastly, using the three features, AcrNET (as given in Fig. 3(A), (B)) is usually better than the shallow learning models (as provided in Table 3 and Table 4), which suggests the usefulness of the three features and the effectiveness of deep learning methods for this problem.

Besides, we also conducted comparison experiments between CNN, RNN and LSTM. We used RNN on varying lengths features, i.e. secondary structure, solvent accessibility and sequence. Then we concatenated the results from RNN with evolutionary features and Transformer features to DNN. We used the same structure in LSTM experiment as RNN. We applied two layers RNN and LSTM with 64 hidden state. Table 16 shows the results of five-fold crossvalidation test. From the result CNN is slightly better than RNN, whereas LSTM might be too sophisticated for these well-defined features.

**Table 16:**
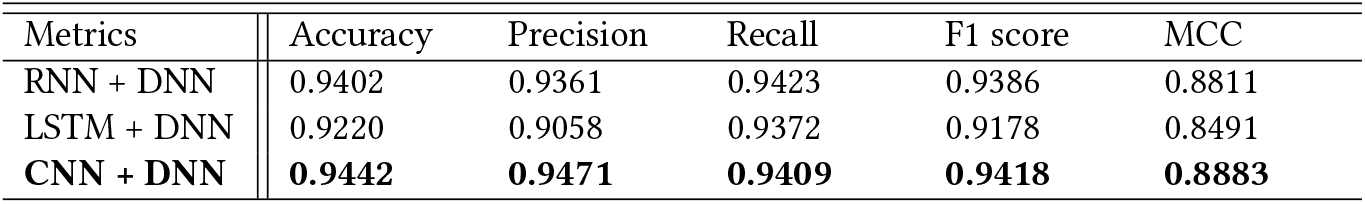
Model influence on the prediction results. We compare the performance between RNN, LSTM and CNN. We used RNN and LSTM along the features with varying lengths, which is the same features that we used CNN on. And then injected the results from RNN or LSTM to DNN.

| Metrics | Accuracy | Precision | Recall | F1 score | MCC |
| --- | --- | --- | --- | --- | --- |
| RNN + DNN | 0.9402 | 0.9361 | 0.9423 | 0.9386 | 0.8811 |
| LSTM + DNN | 0.9220 | 0.9058 | 0.9372 | 0.9178 | 0.8491 |
| **CNN + DNN** | **0.9442** | **0.9471** | **0.9409** | **0.9418** | **0.8883** |

### Classification performance on Acr classes

To further estimate the capacity of our model in predicting the specific class that an Acr protein belongs to, we utilize the same features in the classification task as described in **Extracting evolutionary features and structure features for comprehensive protein representation.**. The comparison of classification problem on Acr categories and corresponding ablation studies are demonstrated in Table 17 and Fig. 3(D) respectively. We can observe from the histograms in Fig. 3(D) that the prediction results obtained using each type of feature are similar, which indicates that the Acr classification task is relatively straightforward when using these inputs. The evolutionary features obtained from PSSM can achieve slightly better results than other features in both single feature and combination with others. The graph may reveal some correlations between Acr classes and their features. To compare AcrNET with the baseline, we adopt the one-vs-rest strategy to AcRanker and PaCRISPR to convert the binary prediction methods to multi-class prediction methods. Because Acrs belonging to the same class are more likely to form motif sequences, Hidden Markov Model (HMM) may also be able to capture these motif features. Thus, we apply HMM on protein sequences and use it as a baseline for Acr classification comparison. AcrNET outperforms these methods across all the metrics, especially on macro-average metrics (improved by around 20% regarding F1 score), suggesting that AcrNET is an unbiased predictor, performing well on the rare Acr classes. Such accurate, detailed predictions from our model can facilitate biological experiments and have the potential of inspiring more insightful studies on the working mechanisms of Acr proteins. In addition to separated prediction and classification problems, we also examined non-Acr samples as the sixth class, and do classification on all data, which means that we combine prediction and classification together. The results in Table 18 and Table 19 show that separate prediction and classification have better results.

**Table 17:** Detailed class prediction performance comparison. We adopt the one-vs-rest strategy for AcRanker and PaCRISPR, converting binary classification methods to five-class classification methods, and compare their performance on the class prediction problem with AcrNET (“mi”:micro-average, “ma”: macro-average). AcrNET outperforms the other methods across all the evaluation criteria significantly and consistently, especially on macro-average, suggesting that AcrNET is an unbiased predictor for small classes. Results in this table are averaged over 10 different random seeds in our experiments.

|  | Accuracy (mi) | Accuracy (ma) | Precision (ma) | Recall (ma) | F1 score (ma) |
| --- | --- | --- | --- | --- | --- |
| AcRanker | 0.8903 | 0.6318 | 0.8532 | 0.6318 | 0.6830 |
| PaCRISPR | 0.8903 | 0.5552 | 0.7071 | 0.5552 | 0.5911 |
| HMM | 0.8600 | 0.4994 | 0.7755 | 0.4994 | 0.5607 |
| **AcrNET** | **0.9480** | **0.8583** | **0.8676** | **0.8583** | **0.8531** |

**Table 18:** Each class prediction performance in AcrNET. We compare classification performance of each class in AcrNET.

| Classes | Accuracy | Precision | Recall | F1 score |
| --- | --- | --- | --- | --- |
| II-A | 0.9801 | 0.9875 | 0.9802 | 0.9836 |
| I-F | 0.9199 | 0.8414 | 0.9199 | 0.8768 |
| I-D | 0.9889 | 0.9567 | 0.9889 | 0.9710 |
| II-C | 0.7915 | 0.7892 | 0.7916 | 0.7646 |
| others | 0.6110 | 0.7630 | 0.6110 | 0.6696 |

**Table 19:** Use non-Acr as the sixth class in classification. We add non-Acr samples as the sixth class and classify all data together. In this case we can solve prediction and classification problem at the same time. Whereas the performance is not as good as separate prediction and classification.

| Classes | Accuracy | Precision | Recall | F1 score |
| --- | --- | --- | --- | --- |
| II-A | 0.9690 | 0.9836 | 0.9690 | 0.9761 |
| I-F | 0.8770 | 0.7848 | 0.8770 | 0.8148 |
| I-D | 0.9243 | 0.8938 | 0.9243 | 0.8999 |
| II-C | 0.7279 | 0.8475 | 0.7279 | 0.7622 |
| others | 0.3386 | 0.5489 | 0.3386 | 0.3932 |
| non-Acr | 0.9554 | 0.9483 | 0.9554 | 0.9516 |
| Macro | 0.7987 | 0.8344 | 0.7987 | 0.7996 |
| Weighted | 0.9372 | 0.9399 | 0.9372 | 0.9356 |

### AcrNET learns Acr motifs implicitly

To explain the hidden rule of model prediction, we conduct motif analysis to study the Acr sequence and structure patterns. MEME [1] can find motif sequences in the protein data. Motif analysis tool can be found at https://meme-suite.org/meme/tools/meme. We used MEME under Motif Discovery. Input the protein sequence file and keep other default value to get the results. The results provide overall motif sequences and motif sequences for each protein. Applying MEME to the Acr dataset, we show that Acrs belonging to the same class are more likely to form motif sequences. To further investigate the structure pattern of these motif sequences, we utilize an accurate protein structure prediction method, AlphaFold. Fig. 4 presents some motif sequence and structure results. Such structures reveal that these motif sequences correspond to highly similar protein secondary structures, which may serve as the hidden rule for AcrNET. Also, replacing motif sequences will change these important protein secondary structures, which may make positive samples lose Acr-related features. To verify this conjecture, we conduct motif mutation by replacing the motif sequences with random sequences for randomly selected Acrs, including Acr0498 (AcrFI11), Acr0562 (AcrIIA7), Acr0559 (AcrIIA8), and Acr0560 (AcrIIA9). The native Acr sequences and mutated sequences are inputted to AcrNET for Acr prediction. The results show that the native Acrs are all predicted as Anti-CRISPR successfully, and all the mutated sequences are predicted as non-anti-CRISPR. Table 20 and 21 show the prediction results. The above results suggest that AcrNET learns the important motifs of the Acr sequences implicitly, which serves as the foundation of its prediction. Considering rigorousness, we also mutate the non-motif sequences and follow the same steps mentioned above to study their effects. The results from Table 22 show that three mutated sequences are still predicted as anti-CRISPR with only one exception, suggesting that our model indeed captures the important motif information in most Acr sequences.

**Figure 4:**
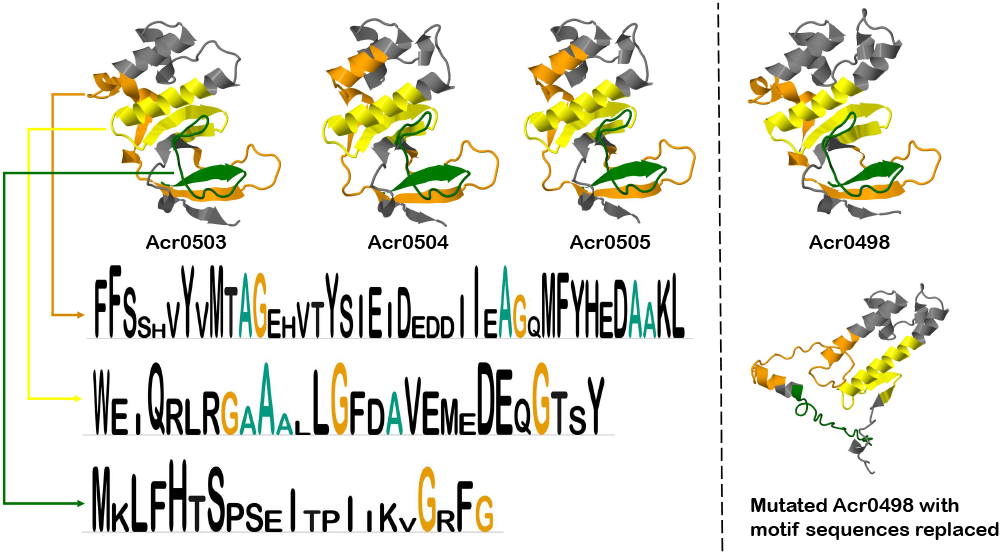
Motif analysis results. The structures of four samples Acr0503, Acr0504, Acr0505, and Acr0498 from the same type AcrFI11 are presented. Three found motifs are listed, and the corresponding structures are colored orange, yellow and green. The lengths of three motif sequences are 41, 28 and 21. The structure of mutated Acr0498, whose motif sequence is replaced by random sequences, is also shown.

**Table 20:** AcrNET prediction confidence scores of native Acr sequences.

|  | Acr0498 | Acr0562 | Acr0559 | Acr0560 |
| --- | --- | --- | --- | --- |
| Acr | 1 | 0.9881 | 1 | 1 |
| Non-Acr | 0 | 0.0119 | 0 | 0 |

**Table 21:** AcrNET prediction confidence scores of mutated Acr sequences with motif sequences replaced by random sequences.

|  | Acr0498 | Acr0562 | Acr0559 | Acr0560 |
| --- | --- | --- | --- | --- |
| Acr | 0 | 0.0015 | 0.0001 | 0.1105 |
| Non-Acr | 1 | 0.9985 | 0.9999 | 0.8895 |

**Table 22:** AcrNET prediction confidence scores of mutated Acr sequences with non-motif sequences replaced by random sequences.

|  | Acr0498 | Acr0562 | Acr0559 | Acr0560 |
| --- | --- | --- | --- | --- |
| Acr | 0.0922 | 1 | 1 | 1 |
| Non-Acr | 0.9078 | 0 | 0 | 0 |

### Docking methods validate our predictions

Biological experimental validation is time-consuming and expensive, and we wish to facilitate it with protein-protein docking and perform more comprehensive prediction results. For a new candidate Acr protein, we predict protein structure with AlphaFold and investigate the interaction between the protein and its target using protein-protein docking tools, which could also provide information about the Acr mechanism. We used the official implementation of AlphaFold v2.0 to perform the protein structure prediction. The package is from: https://github.com/deepmind/alphafold. Following the tutorial, we setup AlphaFold by downloading the genetic databases and model parameters after building the docker environment on our Linux machine. Then AlphaFold can predict the structures for the Acr proteins by taking the sequence FASTA files as input. Here is a slightly simplified version of AlphaFold on Colab Notebook: https://colab.research.google.com/github/deepmind/alphafold/blob/main/notebooks/AlphaFold.ipynb, which is easier to run. As for docking tools, various tools are available including ClusPro [6, 22, 23, 37], HDOCK [15, 16, 42–44], LzerD [5] and Schrödinger [47]. We study the interaction between Acr0275 from Anti-CRISPRdb and its receptor by HDOCK. We mutated the motif sequence from the Acr with random sequences and conducted docking for both the native protein and the mutated one. The docking results are shown in Fig. 5. The docking energy scores increased from −364.51 (predicted as Acr by AcrNET) to −254.79 (predicted as non-Acr by AcrNET), showing the mutated proteins has less stable interaction with the original receptor. Understandably, the current docking method may not be 100% accurate, but it could facilitate the discovery and study of Acr proteins, together with AcrNET and AlphaFold. Further improvement on the tools could lead to better toolkits for Acr investigation.

**Figure 5:**
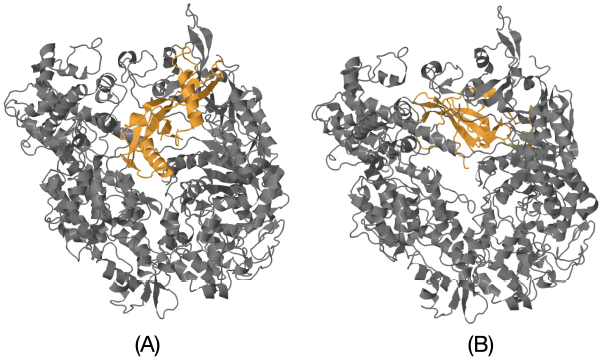
Acr0275 and receptor docking results with HDOCK. We check the docking results of both the native Acr0275 structure and the mutated one, with the motif sequence replaced and its structure predicted by AlphaFold. The structure in orange is the Acr sample, and the structure in grey is the receptor. (A) Docking result of the native Acr0275 and its receptor. (B) Docking result of the mutated Acr0275 with motif sequences replaced and its receptor.

## 3 CONCLUSION

Performing accurate anti-CRISPR predictions can help us reduce off-target accidents in gene editing and develop phage therapy. Here, we propose a deep learning method, AcrNET, for anti-CRISPR prediction. When receiving a protein sequence as input, AcrNET will accurately estimate whether this sequence is an Acr and further predict its specific type if it is an Acr, assisting biological identification of new Acrs efficiently. Without any prior knowledge, using biological experiments to verify whether a protein sequence is an Acr and its specific category is often very time consuming and expensive, which is nearly impossible with the rapid emergence of a large number of protein sequences. AcrNET overcomes the limitation of the experimental methods that rely heavily on the reaction between CRISPR-Cas and Acr to find new Acrs, thus can find Acrs from the large-scale protein database. Furthermore, the proposed AcrNET can carry out accurate prediction and classification tasks efficiently, which is very important for practical applications in big protein databases. By predicting Acr accurately, AcrNET provides useful prior knowledge for biological researchers to identify Acr more efficiently. To further help the community, we provide an online web-server https://proj.cse.cuhk.edu.hk/aihlab/AcrNET/. Due to limited computing power, we only support input files containing no more than three sequences whose length is less than 300. For higher computing needs, please download our program from https://github.com/banma12956/AcrNET and run it locally.

AcrNET achieves high accuracy on the Acr prediction tasks. One important reason is that we introduce the Transformer learning algorithm to deal with the problem of data scarcity. Previous methods only study small sizes of Acr datasets, which makes it difficult to learn general and informative patterns. As a result, these models cannot perform accurate Acr predictions and provide limited prior knowledge on protein sequences, which still need a large number of biological experiments for verification, causing an insufficient driving force for practical usage. By introducing the Transformer module, we can successfully generate effective representations by considering the structural information of protein sequences themselves and using the knowledge of the common properties of protein sequences learned from extensive databases in the pre-training process. These informative representations promote the training of the model effectively and improve the final prediction results, which will greatly decrease the time and cost of biological verification. Such an idea and features can also be applied to similar computational problems with limited annotated data.

The proposed AcrNET method provides the first computational solution for predicting Acr classes, which have not been studied before. Unlike other methods, which rely on the CRISPR-Cas system to perform predictions, the proposed AcrNET can directly predict the specific types of Acr without such limitation. Experimental results demonstrate that the four evolutionary features result in better prediction performance than other features, indicating that the specific Acr types are more closely related to the protein evolutionary information.

The motif analysis shows the importance of motif sequences and their corresponding structures to Acrs prediction. Acrs belonging to the same category are more likely to have similar motif sequences, and these motif sequences correspond to highly similar protein secondary structures, which are the unique features of this type of Acrs. Such sequence patterns and structure patterns can be learned by AcrNET implicitly and serve as the prediction factors. Experiment results show that Acrs with motif sequences replaced will be predicted as non-Acrs by AcrNET, which indicates the prediction basis of AcrNET.

The success of AlphaFold and protein-protein docking analysis enhance our analysis pipeline. We can simulate the interaction between Acr and CRISPR-Cas proteins by utilizing docking tools before biological experiments. These tools can provide useful information, including docking position and docking energy, which can assist in designing and implementing biological experiments.

Despite the great improvement of AcrNET over the previous methods, our method could be improved further with an even larger dataset. Also, information representation of the protein structure from AlphaFold could boost our model, although currently, it is still time-consuming to run AlphaFold, and efficient protein 3D structure representation is still under exploration. Finally, with the assistance of AlphaFold and docking tools, we can illustrate the Acr mechanism to some extent. However, they are not within the AcrNET model. In the future, it will be desirable to design a comprehensive deep learning model, which can perform Acr prediction and illustrate its mechanisms simultaneously.

## REFERENCES

[1] Timothy L Bailey, Charles Elkan, et al. 1994. Fitting a mixture model by expectation maximization to discover motifs in bipolymers. (1994).

[2] Amos Bairoch, Rolf Apweiler, Cathy H. Wu, Winona C. Barker, Brigitte Boeckmann, Serenella Ferro, Elisabeth Gasteiger, Hongzhan Huang, Rodrigo Lopez, Michele Magrane, Maria Jesus Martin, Darren A. Natale, Claire O’Donovan, Nicole Redaschi, and Lai-Su L. Yeh. 2007. The Universal Protein Resource (UniProt). Nucleic Acids Research 35 (2007), D193 – D197.

[3] Joe Bondy-Denomy, April Pawluk, Karen L Maxwell, and Alan R Davidson. 2013. Bacteriophage genes that inactivate the CRISPR/Cas bacterial immune system. Nature 493, 7432 (2013), 429–432.

[4] Shenyang Chen, Qingxiong Tan, Jingchen Li, and Yu Li. 2021. USPNet: unbiased organism-agnostic signal peptide predictor with deep protein language model. bioRxiv (2021).

[5] Charles Christoffer, Siyang Chen, Vijay Bharadwaj, Tunde Aderinwale, Vidhur Kumar, Matin Hormati, and Daisuke Kihara. 2021. LZerD webserver for pairwise and multiple protein–protein docking. Nucleic Acids Research (2021).

[6] Israel T Desta, Kathryn A Porter, Bing Xia, Dima Kozakov, and Sandor Vajda. 2020. Performance and its limits in rigid body protein-protein docking. Structure 28, 9 (2020), 1071–1081.

[7] Jacob Devlin, Ming-Wei Chang, Kenton Lee, and Kristina Toutanova. 2019. BERT: Pre-training of Deep Bidirectional Transformers for Language Understanding. ArXiv abs/1810.04805 (2019).

[8] Shuyan Ding, Yan Li, Zhuoxing Shi, and Shoujiang Yan. 2014. A protein structural classes prediction method based on predicted secondary structure and PSI-BLAST profile. Biochimie 97 (2014), 60–65.

[9] Chuan Dong, Ge-Fei Hao, Hong-Li Hua, Shuo Liu, Abraham Alemayehu Labena, Guoshi Chai, Jian Huang, Nini Rao, and Feng-Biao Guo. 2018. Anti-CRISPRdb: a comprehensive online resource for anti-CRISPR proteins. Nucleic acids research 46, D1 (2018), D393–D398.

[10] Chuan Dong, Dong-Kai Pu, Cong Ma, Xin Wang, Qing-Feng Wen, Zhi Zeng, and Feng-Biao Guo. 2020. Precise detection of Acrs in prokaryotes using only six features. bioRxiv (2020).

[11] Qiwen Dong, Shuigeng Zhou, and Jihong Guan. 2009. A new taxonomy-based protein fold recognition approach based on autocross-covariance transformation. Bioinformatics 25, 20 (2009), 2655–2662.

[12] Simon Eitzinger, Amina Asif, Kyle E Watters, Anthony T Iavarone, Gavin J Knott, Jennifer A Doudna, and Fayyaz ul Amir Afsar Minhas. 2020. Machine learning predicts new anti-CRISPR proteins. Nucleic acids research 48, 9 (2020), 4698–4708.

[13] Ayal B Gussow, Allyson E Park, Adair L Borges, Sergey A Shmakov, Kira S Makarova, Yuri I Wolf, Joseph Bondy-Denomy, and Eugene V Koonin. 2020. Machine-learning approach expands the repertoire of anti-CRISPR protein families. Nature communications 11, 1 (2020), 1–12.

[14] Geoffrey Hinton, Simon Osindero, Max Welling, and Yee-Whye Teh. 2006. Unsupervised discovery of nonlinear structure using contrastive backpropagation. Cognitive science 30, 4 (2006), 725–731.

[15] Sheng-You Huang and Xiaoqin Zou. 2008. An iterative knowledge-based scoring function for protein–protein recognition. Proteins: Structure, Function, and Bioinformatics 72, 2 (2008), 557–579.

[16] Sheng-You Huang and Xiaoqin Zou. 2014. A knowledge-based scoring function for protein-RNA interactions derived from a statistical mechanics-based iterative method. Nucleic acids research 42, 7 (2014), e55–e55.

[17] Alexander P Hynes, Geneviève M Rousseau, Marie-Laurence Lemay, Philippe Horvath, Dennis A Romero, Christophe Fremaux, and Sylvain Moineau. 2017. An anti-CRISPR from a virulent streptococcal phage inhibits Streptococcus pyogenes Cas9. Nature microbiology 2, 10 (2017), 1374–1380.

[18] Ganesh S. Jedhe and Paramjit S. Arora. 2021. Chapter One - Hydrogen bond surrogate helices as minimal mimics of protein -helices. In Synthetic and Enzymatic Modifications of the Peptide Backbone, E. James Petersson (Ed.). Methods in Enzymology, Vol. 656. Academic Press, 1–25. https://doi.org/10.1016/bs.mie.2021.04.007

[19] Wolfgang Kabsch and Christian Sander. 1983. Dictionary of protein secondary structure: pattern recognition of hydrogen-bonded and geometrical features. Biopolymers: Original Research on Biomolecules 22, 12 (1983), 2577–2637.

[20] Morten Källberg, Haipeng Wang, Sheng Wang, Jian Peng, Zhiyong Wang, Hui Lu, and Jinbo Xu. 2012. Template-based protein structure modeling using the RaptorX web server. Nature protocols 7, 8 (2012), 1511–1522.

[21] Eugene V Koonin, Kira S Makarova, and Feng Zhang. 2017. Diversity, classification and evolution of CRISPR-Cas systems. Current opinion in microbiology 37 (2017), 67–78.

[22] Dima Kozakov, Dmitri Beglov, Tanggis Bohnuud, Scott E Mottarella, Bing Xia, David R Hall, and Sandor Vajda. 2013. How good is automated protein docking? Proteins: Structure, Function, and Bioinformatics 81, 12 (2013), 2159–2166.

[23] Dima Kozakov, David R Hall, Bing Xia, Kathryn A Porter, Dzmitry Padhorny, Christine Yueh, Dmitri Beglov, and Sandor Vajda. 2017. The ClusPro web server for protein–protein docking. Nature protocols 12, 2 (2017), 255–278.

[24] Yu Li, Sheng Wang, Ramzan Umarov, Bingqing Xie, Ming Fan, Lihua Li, and Xin Gao. 2018. DEEPre: sequence-based enzyme EC number prediction by deep learning. Bioinformatics 34, 5 (2018), 760–769.

[25] Taigang Liu, Xiaoqi Zheng, and Jun Wang. 2010. Prediction of protein structural class for low-similarity sequences using support vector machine and PSI-BLAST profile. Biochimie 92, 10 (2010), 1330–1334.

[26] Nicole D Marino, Rafael Pinilla-Redondo, Bálint Csörgő, and Joseph Bondy-Denomy. 2020. Anti-CRISPR protein applications: natural brakes for CRISPR-Cas technologies. Nature methods 17, 5 (2020), 471–479.

[27] Linus Pauling, Robert B Corey, and Herman R Branson. 1951. The structure of proteins: two hydrogen-bonded helical configurations of the polypeptide chain. Proceedings of the National Academy of Sciences 37, 4 (1951), 205–211.

[28] April Pawluk, Alan R Davidson, and Karen L Maxwell. 2018. Anti-CRISPR: discovery, mechanism and function. Nature Reviews Microbiology 16, 1 (2018), 12–17.

[29] April Pawluk, Raymond HJ Staals, Corinda Taylor, Bridget NJ Watson, Senjuti Saha, Peter C Fineran, Karen L Maxwell, and Alan R Davidson. 2016. Inactivation of CRISPR-Cas systems by anti-CRISPR proteins in diverse bacterial species. Nature microbiology 1, 8 (2016), 1–6.

[30] Alec Radford, Jeff Wu, Rewon Child, David Luan, Dario Amodei, and Ilya Sutskever. 2019. Language Models are Unsupervised Multitask Learners.

[31] Roshan Rao, Joshua Meier, Tom Sercu, Sergey Ovchinnikov, and Alexander Rives. 2021. Transformer protein language models are unsupervised structure learners. bioRxiv (2021).

[32] Benjamin J Rauch, Melanie R Silvis, Judd F Hultquist, Christopher S Waters, Michael J McGregor, Nevan J Krogan, and Joseph Bondy-Denomy. 2017. Inhibition of CRISPR-Cas9 with bacteriophage proteins. Cell 168, 1-2 (2017), 150–158.

[33] Alexander Rives, Joshua Meier, Tom Sercu, Siddharth Goyal, Zeming Lin, Jason Liu, Demi Guo, Myle Ott, C Lawrence Zitnick, Jerry Ma, et al. 2021. Biological structure and function emerge from scaling unsupervised learning to 250 million protein sequences. Proceedings of the National Academy of Sciences 118, 15 (2021).

[34] Samuel Sledzieski, Rohit Singh, Lenore Cowen, and Bonnie Berger. 2021. Sequence-based prediction of protein-protein interactions: a structure-aware interpretable deep learning model. bioRxiv (2021).

[35] Sabrina Y Stanley and Karen L Maxwell. 2018. Phage-encoded anti-CRISPR defenses. Annual review of genetics 52 (2018), 445–464.

[36] Baris E Suzek, Yuqi Wang, Hongzhan Huang, Peter B McGarvey, Cathy H Wu, and UniProt Consortium. 2015. UniRef clusters: a comprehensive and scalable alternative for improving sequence similarity searches. Bioinformatics 31, 6 (2015), 926–932.

[37] Sandor Vajda, Christine Yueh, Dmitri Beglov, Tanggis Bohnuud, Scott E Mottarella, Bing Xia, David R Hall, and Dima Kozakov. 2017. New additions to the C lus P ro server motivated by CAPRI. Proteins: Structure, Function, and Bioinformatics 85, 3 (2017), 435–444.

[38] Katharina G. Wandera, Omer S. Alkhnbashi, Harris v.I. Bassett, Alexander Mitrofanov, Sven Hauns, Anzhela Migur, Rolf Backofen, and Chase L. Beisel. 2022. Anti-CRISPR prediction using deep learning reveals an inhibitor of Cas13b nucleases. Molecular cell (2022).

[39] Jiawei Wang, Wei Dai, Jiahui Li, Qi Li, Ruopeng Xie, Yanju Zhang, Christopher Stubenrauch, and Trevor Lithgow. 2021. AcrHub: an integrative hub for investigating, predicting and mapping anti-CRISPR proteins. Nucleic Acids Research 49, D1 (2021), D630–D638.

[40] Jiawei Wang, Wei Dai, Jiahui Li, Ruopeng Xie, Rhys A Dunstan, Christopher Stubenrauch, Yanju Zhang, and Trevor Lithgow. 2020. PaCRISPR: a server for predicting and visualizing anti-CRISPR proteins. Nucleic acids research 48, W1 (2020), W348–W357.

[41] Jiawei Wang, Bingjiao Yang, Jerico Revote, Andre Leier, Tatiana T Marquez-Lago, Geoffrey Webb, Jiangning Song, Kuo-Chen Chou, and Trevor Lithgow. 2017. POSSUM: a bioinformatics toolkit for generating numerical sequence feature descriptors based on PSSM profiles. Bioinformatics 33, 17 (2017), 2756–2758.

[42] Yumeng Yan, Huanyu Tao, Jiahua He, and Sheng-You Huang. 2020. The HDOCK server for integrated protein–protein docking. Nature protocols 15, 5 (2020), 1829–1852.

[43] Yumeng Yan, Zeyu Wen, Xinxiang Wang, and Sheng-You Huang. 2017. Addressing recent docking challenges: A hybrid strategy to integrate template-based and free protein-protein docking. Proteins: Structure, Function, and Bioinformatics 85, 3 (2017), 497–512.

[44] Yumeng Yan, Di Zhang, Pei Zhou, Botong Li, and Sheng-You Huang. 2017. HDOCK: a web server for protein–protein and protein–DNA/RNA docking based on a hybrid strategy. Nucleic acids research 45, W1 (2017), W365–W373.

[45] Haidong Yi, Le Huang, Bowen Yang, Javi Gomez, Han Zhang, and Yanbin Yin. 2020. AcrFinder: genome mining anti-CRISPR operons in prokaryotes and their viruses. Nucleic acids research 48, W1 (2020), W358–W365.

[46] Qinze Yu, Zhihang Dong, Xingyu Fan, Licheng Zong, and Yu Li. 2021. HMD-AMP: Protein Language-Powered Hierarchical Multi-label Deep Forest for Annotating Antimicrobial Peptides. arXiv preprint arXiv:2111.06023 (2021).

[47] Kai Zhu, Tyler Day, Dora Warshaviak, Colleen Murrett, Richard Friesner, and David Pearlman. 2014. Antibody structure determination using a combination of homology modeling, energy-based refinement, and loop prediction. Proteins: Structure, Function, and Bioinformatics 82, 8 (2014), 1646–1655.

[48] Lingyun Zou, Chonghan Nan, and Fuquan Hu. 2013. Accurate prediction of bacterial type IV secreted effectors using amino acid composition and PSSM profiles. Bioinformatics 29, 24 (2013), 3135–3142.

[49] Zhenzhen Zou, Shuye Tian, Xin Gao, and Yu Li. 2019. mlDEEPre: Multi-functional enzyme function prediction with hierarchical multi-label deep learning. Frontiers in Genetics 9 (2019), 714.

